# The correlation between central and peripheral oxytocin concentrations: a systematic review and meta-analysis

**DOI:** 10.1101/096263

**Authors:** Mathias Valstad, Gail A. Alvares, Maiken Egknud, Anna Maria Matziorinis, Ole A. Andreassen, Lars T. Westlye, Daniel S. Quintana

## Abstract

There is growing interest in the role of the oxytocin system in social cognition and behavior. Peripheral oxytocin concentrations are regularly used to approximate central concentrations in psychiatric research. This methodological approach has obvious appeal given the invasiveness of cerebrospinal fluid collection. However, the validity of this approach and potential moderators of the association between central and peripheral levels are unclear. Thus, we conducted a pre-registered systematic search and meta-analysis of correlations between central and peripheral oxytocin concentrations. A search of databases yielded 17 eligible studies for effect size synthesis and moderator analysis, resulting in a total sample size of 516 participants and subjects. Overall, a positive association between central and peripheral oxytocin concentrations was revealed [r=0.29, 95% CI (0.15, 0.42), p<0.0001], along with a moderate-to-high level of heterogeneity across effect sizes [Q=88.14, p<0.0001], and no evidence of publication bias (p=0.45). This association was significantly moderated by experimental context [Qb(4), p=0.0016]. The strongest association was observed after intranasal oxytocin administration (*r*=0.67, *p*<.0001), a correlation that was significantly greater (*p*=.0002) than the equivalent association under baseline conditions (*r*=0.08, *p*=.31). These results support the use of peripheral levels of oxytocin as a marker of central levels, but only after exogenous oxytocin administration. Despite the popularity of using peripheral OT levels to approximate central levels during baseline conditions, this approach is not supported by the present results.

## Introduction

Oxytocin is a nine amino acid neuropeptide that acts on the widely distributed G-protein coupled oxytocin receptor in humans and almost all other vertebrate species (1). Oxytocin is released both into the central nervous system (CNS) and peripheral circulation from neurosecretory cells in the paraventricular (PVN) and supraoptical (SON) nuclei of the hypothalamus, where most endogenous oxytocin is synthesized. Central and peripheral compartments of the oxytocin system are separated anatomically by the blood-brain barrier, that only in exceptional cases is appreciably permeated by oxytocin (2).

Through central action, oxytocin is critically involved in a range of social behaviors and social cognitive functions (3). Endogenous, or naturally produced, oxytocin appears to co-vary with social cognition at all levels of information processing in humans and other mammals, with similar effects after administration of exogenous oxytocin (4). Growing clinical interest (5) has focused on neurodevelopmental and psychiatric conditions characterized by social cognition and behavioral impairments, such as autism spectrum disorder (ASD) (6,7) and schizophrenia (8), with the hope to explore the potential of oxytocin as a biomarker of these conditions, better understand their potential etiological pathways, and ultimately to ameliorate the associated social-cognitive and behavioral symptoms.

Several methodological approaches have been adopted to the study of oxytocin involvement in normal and impaired social behavior and cognition. These include the measurement of psychological or neurobiological outcomes after administration of exogenous oxytocin, and the assessment of endogenous oxytocin concentration covariance with psychological phenotypes and psychiatric disorder status. While crucial to the latter, concentrations of oxytocin have been sampled within both of these research traditions. Although the social cognitive effects of oxytocin are attributed to central mechanisms, oxytocin concentrations have typically, but not universally, been sampled in peripheral fluids such as blood plasma, saliva, and urine (9). Consequentially, that peripheral oxytocin concentrations approximate central bioavailability of the neuropeptide has been a crucial assumption in research where peripheral oxytocin concentrations are correlated with psychological phenotypes or psychiatric disorder status.

Although some animal research indicates that central release from the hypothalamus and peripheral release via the posterior pituitary is coordinated (10-12), other research does not support this (13,14). Research is also mixed in humans, with some results consistent with related levels of central and peripheral endogenous oxytocin (15), while others report no significant associations (16). After exogenous oxytocin delivered via intranasal administration in humans, one study found a significant association between cerebrospinal fluid (CSF) and blood plasma concentrations of oxytocin (17), while another found no significant association (18). Using peripheral oxytocin concentrations to index central concentrations is clearly appealing, given the more invasive procedures required to collect centrally circulating fluids in humans. However, it is currently unclear whether and when peripheral oxytocin measures can be used to index CNS concentrations and central oxytocin bioavailability.

The present systematic review and meta-analysis synthesized studies in which central and peripheral measures of oxytocin were simultaneously sampled into a summary effect size. The strength of the summary effect size is indicative of the plausibility of peripheral oxytocin as an index for central oxytocin concentrations. As eligible studies were likely to vary in a range of contextual specifications, several potential moderator variables were considered, including experimental paradigm, oxytocin sampling location, subject species, biochemical analysis methods, year of publication, and study quality. Such differences between contexts may contribute to variance in the correlations between central and peripheral oxytocin. Thus, it is possible that peripheral oxytocin can index central oxytocin concentrations in some contexts, but not others. Together, the purpose of this study was to examine whether, and under which circumstances, peripheral oxytocin is a correlate of central oxytocin concentrations.

## Materials and Methods

The systematic search and meta-analysis was conducted in accordance with the PRISMA guidelines (19) (Supplementary material I) and recent recommendations for conducting correlational meta-analyses (20). Prior to the execution of the systematic search and meta-analysis, the protocol for this systematic review and meta-analysis was published (21) and pre-registered on the PROSPERO registry (CRD42015027864).

### Systematic literature search and inclusion of eligible studies

A systematic literature search was performed in two iterations to retrieve studies in which oxytocin had been simultaneously sampled in fluids or tissues located in central (e.g., local extracellular fluid or CSF) or peripheral (e.g. blood plasma or saliva) regions of the body. In the first iteration, a search was performed, using Ovid, in Embase and Medline with the following combination of terms: (oxytocin) AND (concentration^*^ OR level^*^) AND (plasma OR blood OR saliva^*^ OR urin^*^) AND (central OR csf OR “cerebrospinal fluid”). The following constraints were applied to limit search results: the result should be (i) a full-text article or a conference abstract, (ii) written in English, that was (iii) published after 1971, when biochemical analysis of oxytocin content using enzyme immunoassay was made commercially available. The search was conducted on August 2, 2016, and resulted in a total of 572 studies. Out of these, 111 were relevant. A second iteration was performed in which citing articles and reference lists of included studies were examined for remaining relevant studies (Fig. 1). After retrieval, relevant studies were screened for inclusion based on the criterion that effect sizes for the correlation between central and peripheral concentrations of oxytocin must be obtainable. While 121 of the studies retrieved in the systematic search were relevant, only 17 of these satisfied this criterion.

**Figure 1.**
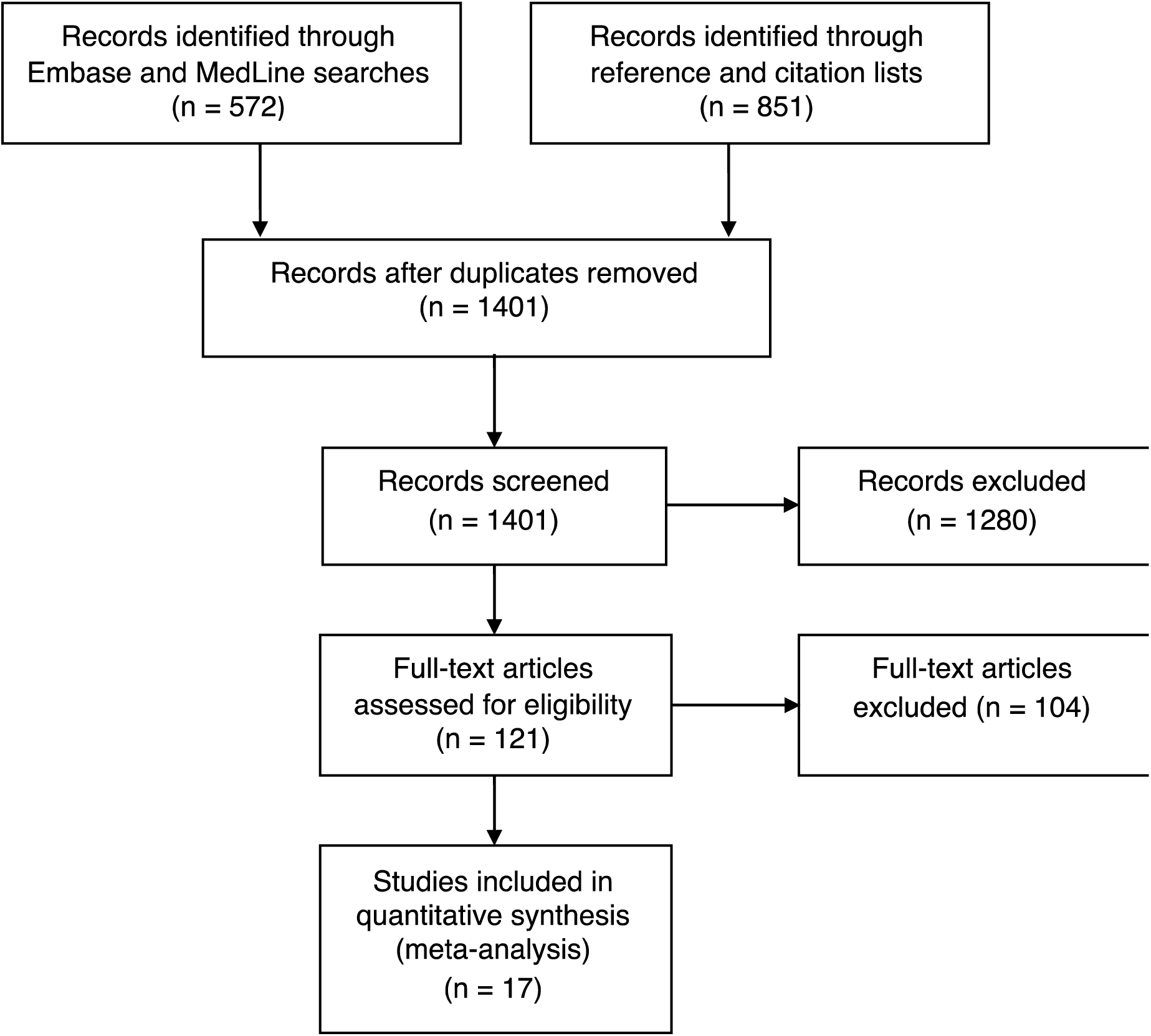
Flowchart of inclusion process. **n**: number of records.

### Data extraction and management

Effect sizes and sample sizes were extracted from eligible studies. For some articles, effect sizes were stated explicitly, or directly obtainable through tables of individual values. In other articles, individual values were represented in graphs such as scatterplots, in which case a web plot digitizer (22) was used for conversion of plots into numerical values. Since some articles contained both a scatterplot and a directly stated effect size, this plot digitizer was validated through comparing effect sizes provided by authors with plot digitizer outputs, revealing almost perfect precision (Supplementary material II). Some articles did not provide relevant effect sizes, individual values in tables, or scatterplots. Since 15 years is a common time frame for the retention of clinical data, authors of such articles published from 2001 were contacted and asked to provide effect sizes. Articles lacking this information that were published before 2001 (n = 69), and studies performed by authors that were not able to respond to the data request (n = 35), were not included in the meta-analysis. Data were extracted from all eligible studies using a custom data extraction form (see Supplementary material III).

### Statistical analysis

Statistical analysis was performed with R statistical software version 3.2.4. (R Core Team, 2016), using the *MAc* (23), *metafor* (24), and *multcomp* (25) R packages. The dataset and script to perform the analyses are available at http://osf/aj55y/

Prior to meta-analytic synthesis, raw effect sizes were transformed to Fischer’s *z* for variance stabilization (26). Raw effect sizes given as Spearman’s ρ were first transformed to Pearson’s *r* according to Gilpin (27), and then transformed to Fischer’s *z* for meta-analysis. For studies reporting several effect sizes, or reporting one effect size based on repeated measures, within-study variance was estimated using the procedure described in supplementary material IV. A random effects model (28), where between-studies variance (τ^2^) was estimated using a restricted maximum likelihood method was used in the synthesis of individual effect sizes into a summary effect size. Outlier diagnostics were also performed to identify potential effect size outliers (24). Point estimates were converted back to Pearson’s *r* for interpretive purposes. The observed variance between studies may be due to heterogeneity (variance in the true effect sizes between studies) and within-study variance. Q, the significance of Q, and I^2^ were computed in order to examine variance and heterogeneity among effect sizes of included studies. I^2^ values of ~25%, ~50%, and ~75% were interpreted as low, moderate, and high, respectively (29).

Potential moderator variables were defined *a priori* (21). Some of the levels for moderator variables were also defined *a priori*, such as the levels *baseline condition* (lack of experimental intervention) and *intranasal administration* for the experimental paradigm moderator. Other levels of moderator variables were adjusted from preplanned analyses *post hoc* based on the specific characteristics of included studies (for details, see Supplementary material V). A random effects model with separate estimates of between studies variance was applied for all categorical moderator variables, yielding summary weighted mean effects and the significance of subgroup effects, which were calculated for each subgroup. Although mammals share essential oxytocin system characteristics, such as production of oxytocin in the hypothalamus, peripheral and central release of oxytocin from hypothalamus, and a blood brain barrier that inhibits diffusion of oxytocin between the CNS and systemic circulation (1), the between-species differences (21) necessitated an additional analysis to examine the role of species in the different effects observed between experimental paradigms. When there were more than two subgroups, pairwise comparisons were performed between all moderator categories with Holm-adjusted p-values to control the family-wise error rate. Meta-regression models were fitted to account for heterogeneity of continuous moderator variables.

### Data quality measures

Small study bias, which includes both publication bias and study quality bias (30)(31), was assessed by visually inspecting a funnel plot and performing Egger’s regression test (24). A significant test (*p* < .05) is indicative of small study bias. A contour enhanced funnel plot, which superimposes key areas of statistical significance (*p* = .1, *p* = .05, *p* = .01), was constructed to specifically assess for risk of publication bias (32). An over-representation of effect sizes in the key areas of significance is indicative of publication bias risk. Since the decision to report a specific effect size, in contrast to the decision to publish a study, is not directly dependent on sample size, the regression test for funnel plot asymmetry does not rule out the possibility that there could be a bias in the type of evidence that is *reported* in published studies. To examine whether this was a source of bias in the set of included studies, the included studies that explicitly stated effect sizes were compared to the studies where effect sizes were obtained by other means, such as data scraping or author request.

There may also be problems with validity of the data that are internal to included studies. A custom risk of bias tool (Supplementary Material VI) was used (by ME and AMM) to systematically assess within-study risk of bias in included studies. This tool was developed by adapting the tool used in another meta-analysis (33) to the context of oxytocin research.

## Results

17 studies yielding 32 effect sizes were included in the meta-analysis (Table 1; Fig. 1). The total number of participants/subjects across studies was 516. Among these, 257 were human, 248 were rodents, 7 were sheep, and 4 subjects were non-human primates.

**Table 1:**
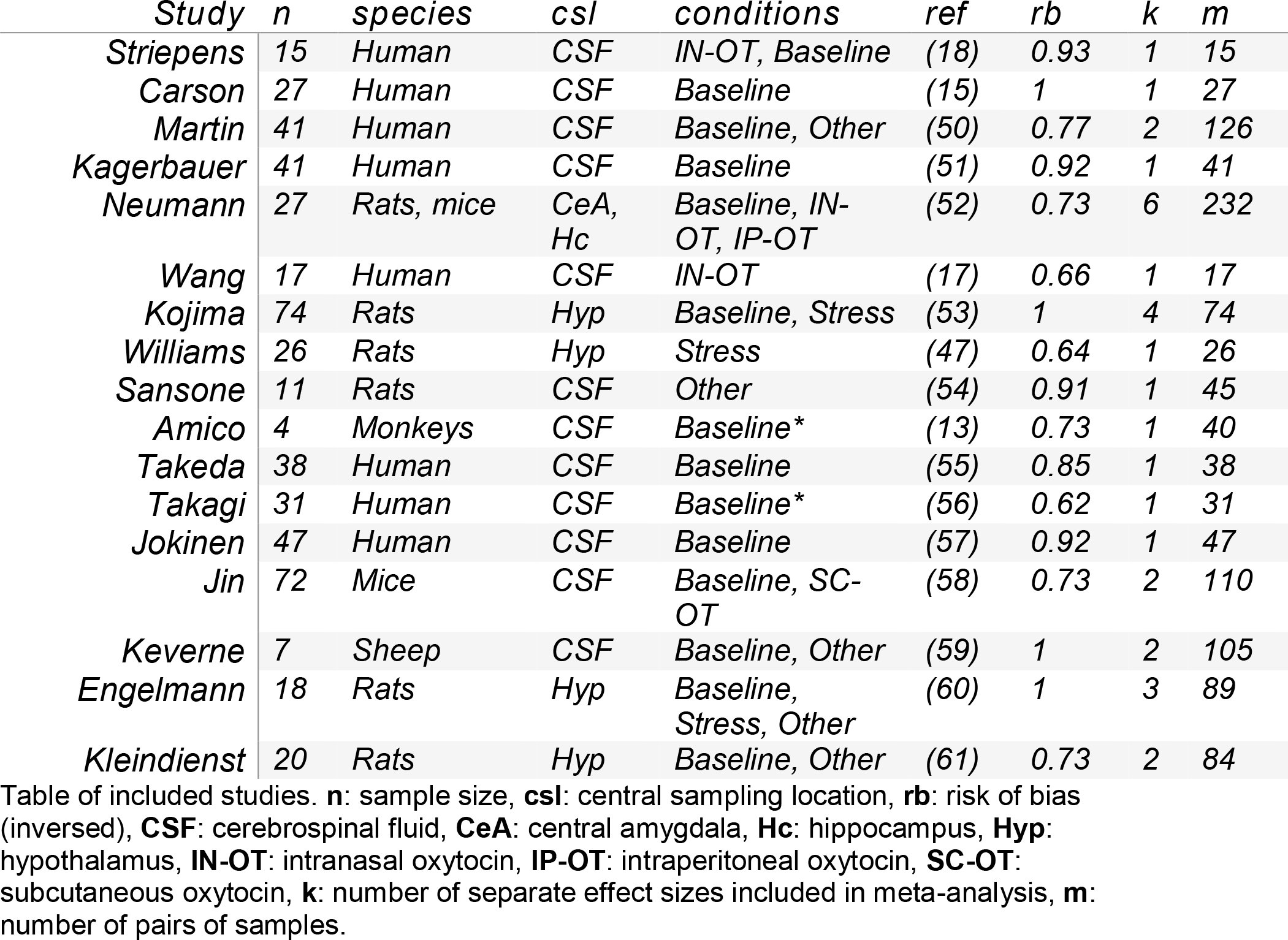
Overview of included studies

### Association between central and peripheral concentrations of oxytocin

There was a positive correlation between central and peripheral concentrations of oxytocin [r = 0.29, 95% CI (0.15, 0.42), *p* < 0.0001; Fig 2]. Egger’s regression test revealed no evidence of publication bias (*p* = .45; Fig 3A). An inspection of the contour enhanced funnel plot did not reveal an over-representation of effect sizes in the significance contours (Fig. 3B). Furthermore, a meta-regression revealed that risk of bias did not influence effect sizes (p= 0.22; Fig 3C). Influence diagnostics identified one potential outlier (17). A sensitivity analysis, which involved re-analysis without the identified outlier, revealed a similar summary effect size as the original analysis that was also statistically significant [r = 0.24, 95% CI (0.12, 0.36), *p* = 0.0002]. As the sensitivity analysis suggested that this single effect size only had a modest effect on the overall meta-analysis, it was retained for the remainder of the analyses. In the total sample of included studies, there was a moderate-to-high level of heterogeneity [Q = 88.14, p < .0001, I^2^ = 63.5% (37.8%, 76.8%)]. Accordingly, moderator analyses were performed to identify sources of heterogeneity.

**Figure 2.**
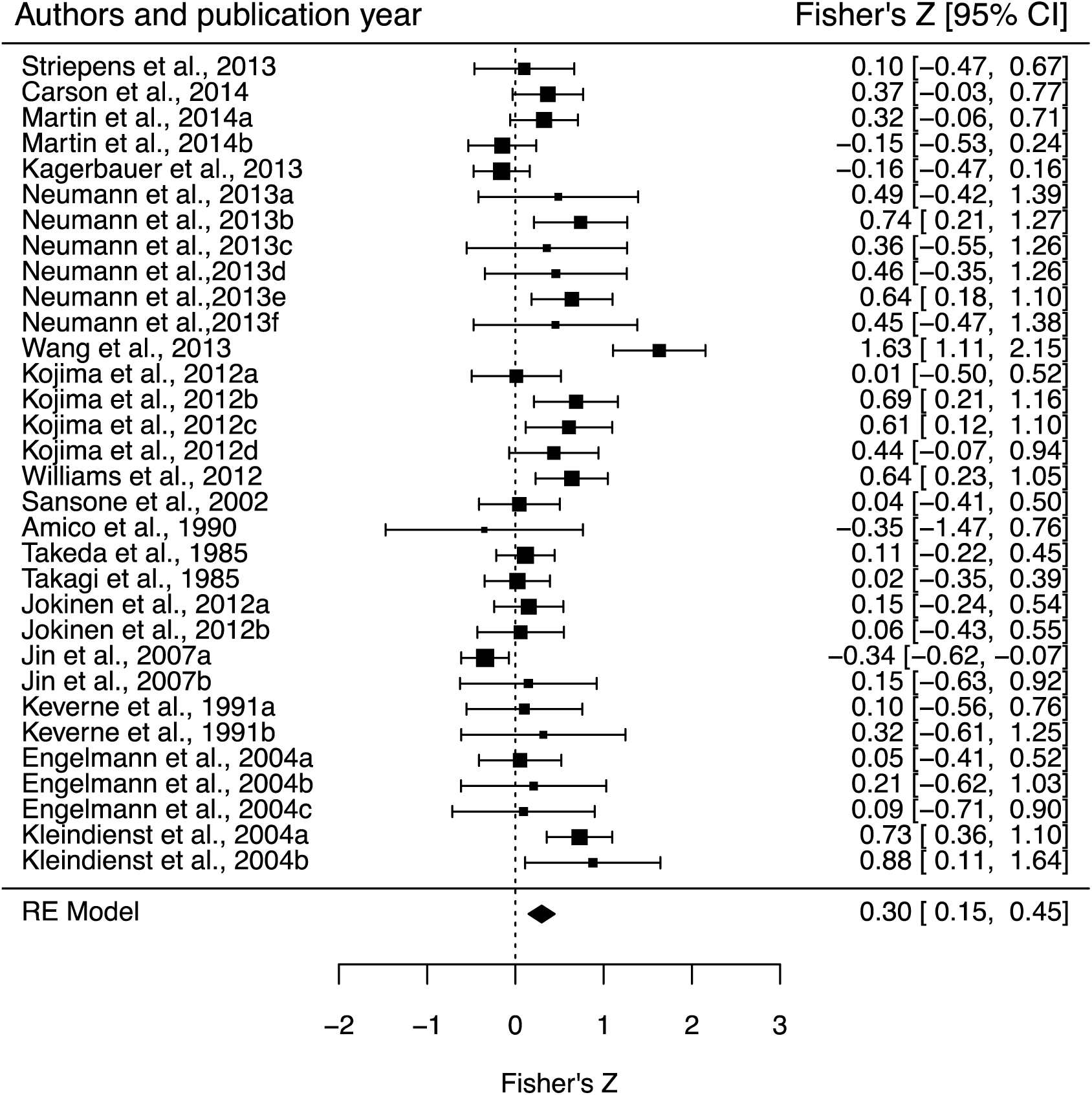
The relationship between peripheral and central oxytocin concentrations. Fisher’s *Z* point estimates are depicted by filled squares, with square sizes indicating the relative weight of each study’s effect size estimate in the analysis. The filled diamond reflects the overall summary effect size [Fisher’s *Z* = 0.30, 95% CI (0.15 to 0.44), *p* < 0.0001]. Error bars and diamond width indicate 95% CIs. Note that the Fisher’s Z summary point estimate slightly differs from the transformed Pearson’s *r* point estimate, which is reported in text. **RE** = Random effects model.

**Figure 3.**
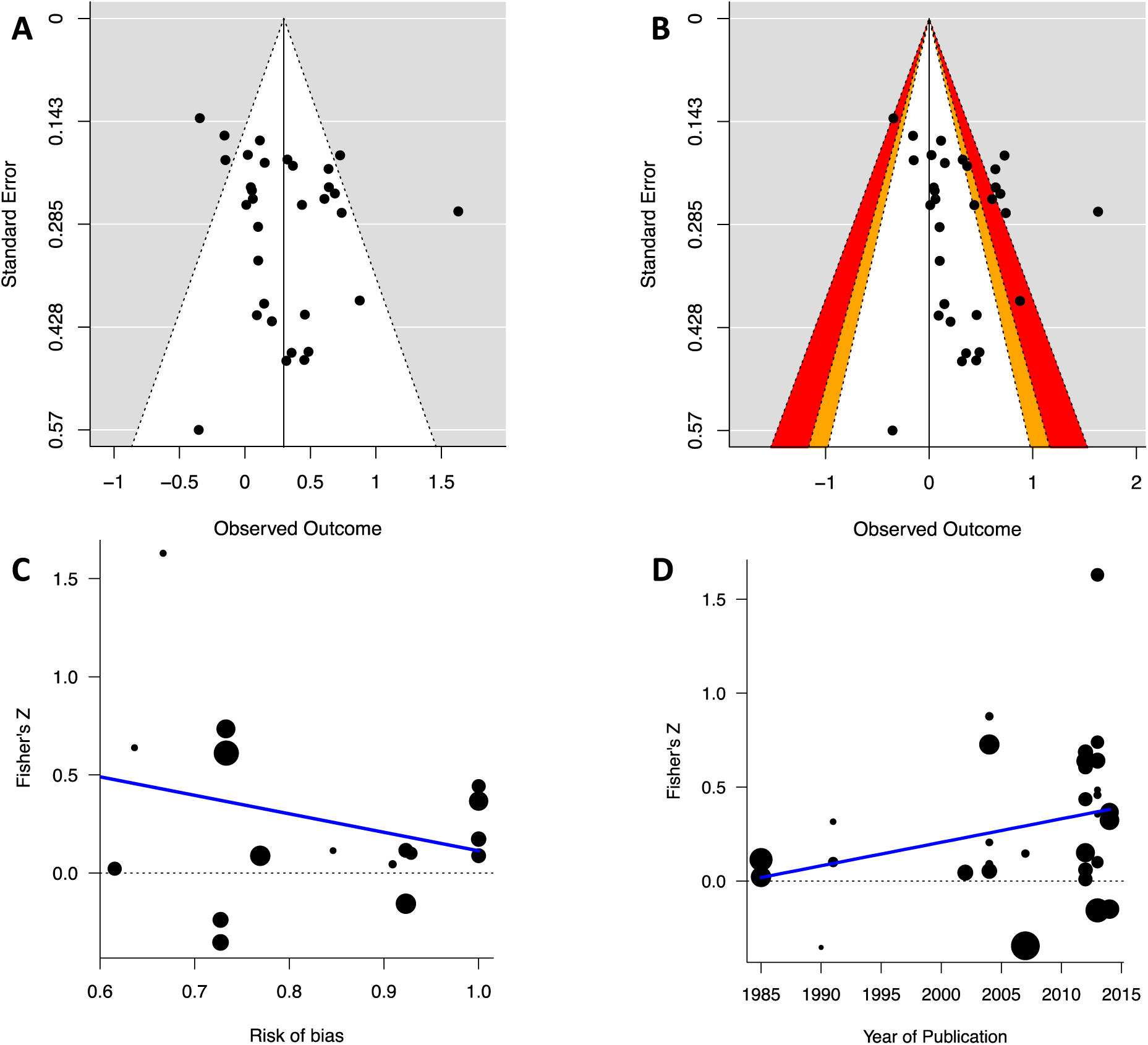
Tests of bias and the impact of moderator variables. Plots A and B illustrate the individual effect sizes on the horizontal axis and corresponding standard errors on the vertical axis. Visual inspection of plots does not reveal evidence of publication bias, as there does not appear to be any asymmetry (A) or an overrepresentation of effect sizes in the orange and red significance contours (B). Meta-regression models revealed no evidence that risk of bias (C; p=0.2204) or publication year (D; *p* = 0.14) had a significant influence on effect size. The solid blue lines in plots C and D represent the respective predicted Fisher’s Z scores based on a mixed-effects models.

### Impact of moderators on effect size

A moderator analysis revealed that part of the heterogeneity in the model was due to the type of experimental paradigm [Q_b_(4) = 17.44, *p* = 0.002; Fig 4A]. Across experimental paradigms, positive associations were observed for the intranasal oxytocin (IN-OT) condition (*r* = .67, p < .0001, k = 4) and after stress interventions (*r* = .49, p = 0.0005, k = 5; Supplementary material VII). In contrast, no association was observed in the baseline condition (r = 0.08, *p* = .31, k = 15). The subgroup effects for the peripheral oxytocin administration category (r = 0.29, *p* = .28, k = 3), as well as for the ‘other’ category (r = 0.30, *p* = .09, k = 5) were not significant. The results for the baseline condition were similar (*r* = 0.10, p=0.26, k = 13) when applying a strict rather than inclusive extension of ‘baseline’. A comparison of all possible pairwise comparisons with Holm corrected *p*-values revealed that the IN-OT point estimate was significantly greater than the baseline point estimate (*p* = .002; Fig. 4A). While there were no other significant pairwise comparisons, the increased stress point estimate compared to baseline point estimate was on the border of statistical significance (*p* = .089). When constrained to human studies, results for the levels of the experimental paradigm moderator were reproduced [Q_b_(2) = 7.56, *p* = 0.02], with no significant correlation in the baseline condition [r = 0.05, 95% CI (-0.19, 0.29), *p* = .58, k = 7, I^2^=0%], and a significant correlation in the intranasal condition [r = 0.71, 95% CI (0.34, 0.89), p = 0.013, k = 2, I^2^ = 93%].

**Figure 4.**
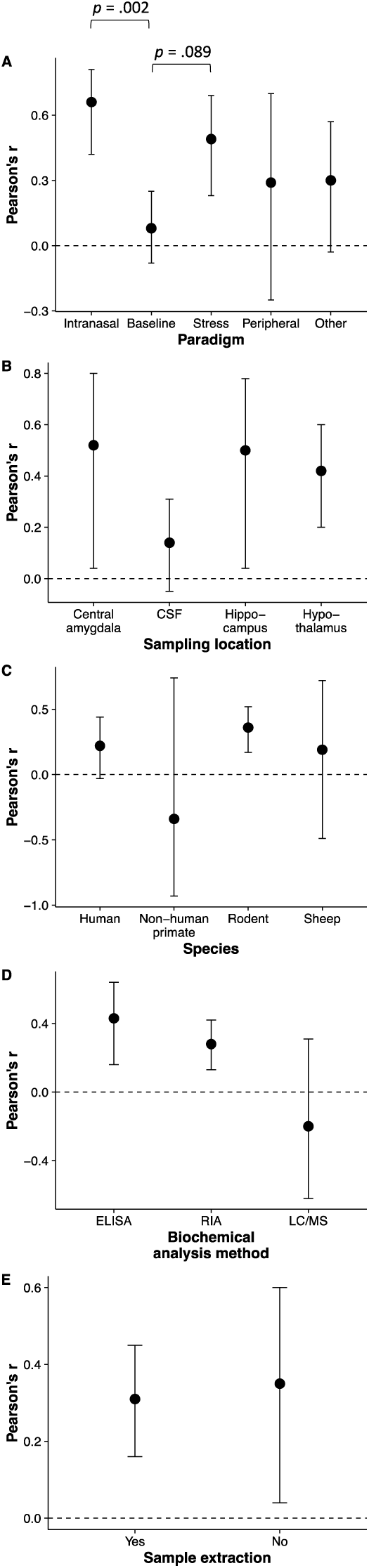
Subgroup effects for the moderators: experimental paradigm (A), central sampling location (B), subject species (C), biochemical analysis method (D), and sample extraction (E). Group summary point estimates are expressed as Pearson’s r with 95% confidence intervals. There was a significant difference in the summary point estimates of endogenous administration and baseline experimental paradigm studies (Holm corrected *p* = .002).

Analysis of the effect of central sampling location on effect sizes was on the border of significance [Q_b_(3) = 6.71, p=0.08; Fig. 4B], suggesting that specific brain sampling location differences may contribute to observed heterogeneity. Across levels of the central sampling location moderator, subgroup effects for hypothalamus (*r* = 0.42, *p* = 0.0003, k = 10), central amygdala (*r* = 0.52, *p* = 0.034, k = 3), and hippocampus (*r* = 0.53, *p* = 0.023, k = 3; Supplementary material VII) were significant. The subgroup effect for samples taken from CSF (*r* = 0.14, *p* = 0.14, k = 16) was not significant. Pairwise comparisons did not reveal any significant difference between any of these subgroups.

The moderator analysis for species was not significant [Q_b_(3) = 1.93, p = 0.59; Fig. 4C], suggesting that species diversity might not contribute to heterogeneity among effect sizes. Across levels of the species moderator, only the subgroup effect for rodents was significant (r = 0.36, *p* = .0004, k = 19; Supplementary material VII). The point estimates for human (*r* = 0.22, *p* = 0.081, k = 10), sheep (r=0.19, *p* = .54, k = 2), and non-human primate (r = -0.34, *p* = .53, k = 1) subgroups were not significant. There was no significant difference between any of the levels of this moderator variable. An exploratory mixed-effect meta-regression model was fitted to assess whether the influence of species type (human vs. rodent) on the correlation between central and peripheral levels varied between experimental paradigms (intranasal oxytocin vs. baseline vs. other). This model revealed no evidence for a significant interaction (Q_b_(2) = 0.43, p =.81).

The biochemical analysis method moderator analysis was not significant [Q_b_(2) = 4.67, p=0.097; Fig. 4D], indicating that this moderator variable is not likely to contribute to heterogeneity among effect sizes. Across the levels of the biochemical analysis method moderator variable, subgroup effects for both RIA (*r* = 0.28, *p* = 0.0005, k = 24) and EIA (*r*=0.43, p=0.002, k = 6) were significant. There was no significant effect for LC/MS (*r* = -0.2, *p* = 0.43, k = 2) and no significant differences between the levels of this moderator variable. The peptide extraction moderator analysis was not significant [Q_b_(1) = 0.05, *p* = 0.82; Fig. 4E]. Both with (*r* = 0.31, *p* = 0.0001, k = 24) and without (*r* = 0.35, *p* = 0.028, k = 6) extraction subgroup effects were significant (Supplementary material VII). Finally, the year of publication did not significantly moderate the relationship between central and peripheral oxytocin concentrations [Q(1) = 2.19, *p* = .14; Fig 3D].

## Discussion

The present systematic meta-analysis revealed a positive correlation between concentrations of oxytocin in blood plasma and oxytocin concentrations in the CNS. However, the association was moderate and showed a high degree of heterogeneity, suggesting that the observed association might not be present across all contexts. Experimental paradigm was the moderator variable most likely to account for this heterogeneity. After IN-OT, as well as after an experimental stressor, there was a positive correlation between central and peripheral oxytocin concentrations. However, in the baseline condition, there was no evidence of correlation, neither for the entire sample of subjects, nor for any of the species analyzed separately. Notably, there was a statistically significant difference between the summary statistic for IN-OT studies and baseline studies. Given the lack of evidence for a correlation between peripheral and central oxytocin levels in the baseline condition, the data suggest blood plasma may not efficiently index central oxytocin concentrations under baseline conditions. Furthermore, this result provides additional indirect evidence for the effectiveness of the blood-brain barrier in restricting oxytocin diffusion between systemic circulation and the CNS (2), as well supporting the hypothesis that under baseline conditions, hypothalamic oxytocin release into blood and into the CNS is uncoordinated (13).

There is a substantial body of research attempting to link peripheral oxytocin concentrations with psychological phenotypes or psychiatric disorder status. Since the social-cognitive effects of oxytocin have been assumed to arise from oxytocin action in the CNS, the assumption that peripheral and central oxytocin concentrations correlate in a baseline condition was crucial in the interpretation of the results from these two approaches (e.g. 34,35). This assumption is called into question by the present data. These results may have two possible, mutually exclusive, implications for the interpretation of studies within these research traditions: either the apparent social cognitive effects are type I errors produced by chance, or the demonstrated covariance between social cognition and endogenous oxytocin in systemic circulation arise from some phenomenon unrelated to central oxytocin levels. The former potential interpretation is consistent with the evidence of publication bias that has surfaced in the field of psychological and psychiatric oxytocin research (9,36,37). The latter interpretation points to a potential peripheral mechanism for the observed social cognitive correlates of basal peripheral oxytocin concentrations. One potential causal mechanism is oxytocin action on peripheral tissues that provide afferent feedback to the CNS (1).

In contrast to what was discovered under baseline conditions, this metaanalysis revealed a positive correlation between central and peripheral oxytocin after intranasal administration of oxytocin. Almost every study examining the effects of exogenous oxytocin on social cognition and behavior in normal and clinical populations have made use of the intranasal delivery route (e.g. 38-41). The motivation behind administering oxytocin intranasally is to obtain non-invasive delivery of oxytocin into the brain. Although vasopressin, which is structurally similar to oxytocin, has been shown to enter the CSF after intranasal administration (42), it is not entirely clear where intranasally administered oxytocin travels, or whether it actually reaches brain areas containing oxytocin receptors such as the hypothalamus or the amygdalae (43). However, recent work in humans comparing intranasal and intravenous oxytocin administration indicates that despite comparable peripheral oxytocin concentrations after both administration routes, social cognitive (44) and neural effects (45) were only observed after intranasal administration. Together, these results are consistent with a direct nose-to-brain transport of intranasally administered oxytocin via olfactory and trigeminal nerve fibers.

In this meta-analysis, a positive association was also found between central and peripheral concentrations of oxytocin after experimental stress induction. Stress induction involved either separation from a mother (46), or a forced swim test (47). As the authors suggest (47), the hypothalamus-pituitary-adrenal (HPA) axis and related hormones such as corticosterone interact with the oxytocin system to regulate stress responses. Such interaction may occur through interneurons between magnocellular and parvocellular neurons in the PVN (48), from which oxytocin and corticotropin-releasing hormone are released. Furthermore, interaction may be mediated through corticosteroid effects on vasoconstriction and heart rate, which in turn could affect oxytocin release through baroreceptors and the vagal feedback system (1,43).

There are some limitations to the study worth mentioning. First, to estimate variances for effect sizes from repeated measures, dependent samples variance estimation was used to control for dependency between samples. Since exact dependencies between repeated samples were unknown, there is a chance that variances for effect sizes obtained in repeated measures designs were slightly overestimated or underestimated, relative to variances for effect sizes obtained in single sample designs. A differential variance estimation would favor one of the two study types with respect to the relative weight they were afforded in the main analysis. However, since there is no a priori reason to believe that study type should impact upon the estimated effect sizes, it is unlikely that this potential bias had any considerable effects on the results. Second, even if there was no evidence for publication bias, or for bias in report of effect sizes, there may be some bias in the subjects sampled for studies where CSF was collected. Across included studies, some of the human participants had medical conditions (15,17,18). Medical conditions are often associated with pain, and pain may influence oxytocin release: in one study, chemical pain stimulation increased oxytocin release within the brain, but not in plasma (49). If pain leads to uncoordinated release, then this may bias the results of this meta-analysis in a negative direction. However, the strongest correlation between central and peripheral concentrations of oxytocin among included studies – which was also identified as a potential outlier – was observed in a sample of headache patients (17). This may point to the opposite possibility that pain could bias the effect sizes of this meta-analysis in a positive direction. To ensure that this study did not inflate the effect size for the IN-OT moderator, a secondary analysis was performed with this study removed, yielding comparable results.

The collection of peripheral oxytocin measures to index central levels has obvious appeal given the difficulties surrounding central collection. However, research has yet to establish whether this is a valid experimental approach. The results of this meta-analysis indicate that there is a positive association between central and peripheral concentrations of oxytocin, but this association depends on experimental context. As there was evidence for a positive association between central and peripheral concentrations of oxytocin after intranasal oxytocin administration and experimental stress induction – both overall and when only including humans in the analysis – blood plasma could be used to approximate central levels under these circumstances. However, as there was no evidence for an association between central and peripheral oxytocin concentrations under baseline conditions, future studies on the role of basal oxytocin in cognition or social behavior should avoid using peripheral oxytocin measures to make inferences on central oxytocin concentrations.

## Acknowledgements

MV received salary support from the Research Council of Norway (RCN) via a grant for students in clinical psychology programmes. GAA is funded by the Cooperative Research Centre for Living with Autism (Autism CRC), established and supported under the Australian Government’s Cooperative Research Centres Program. The Research Council of Norway (RCN) and OptiNose AS contributed to funding this review through a BIA project grant (219483) via salary support to DSQ and project support to OAA, LTW, and DSQ. LTW is supported by the South-Eastern Norway Regional Health Authority (2014097). DSQ is supported by an Excellence Grant from the Novo Nordisk Foundation (NNF16OC0019856). We thank Hege Kristin Ringnes (University of Oslo Library) for providing guidance on our systematic search strategy.

## Supplementary material I: Prisma checklist

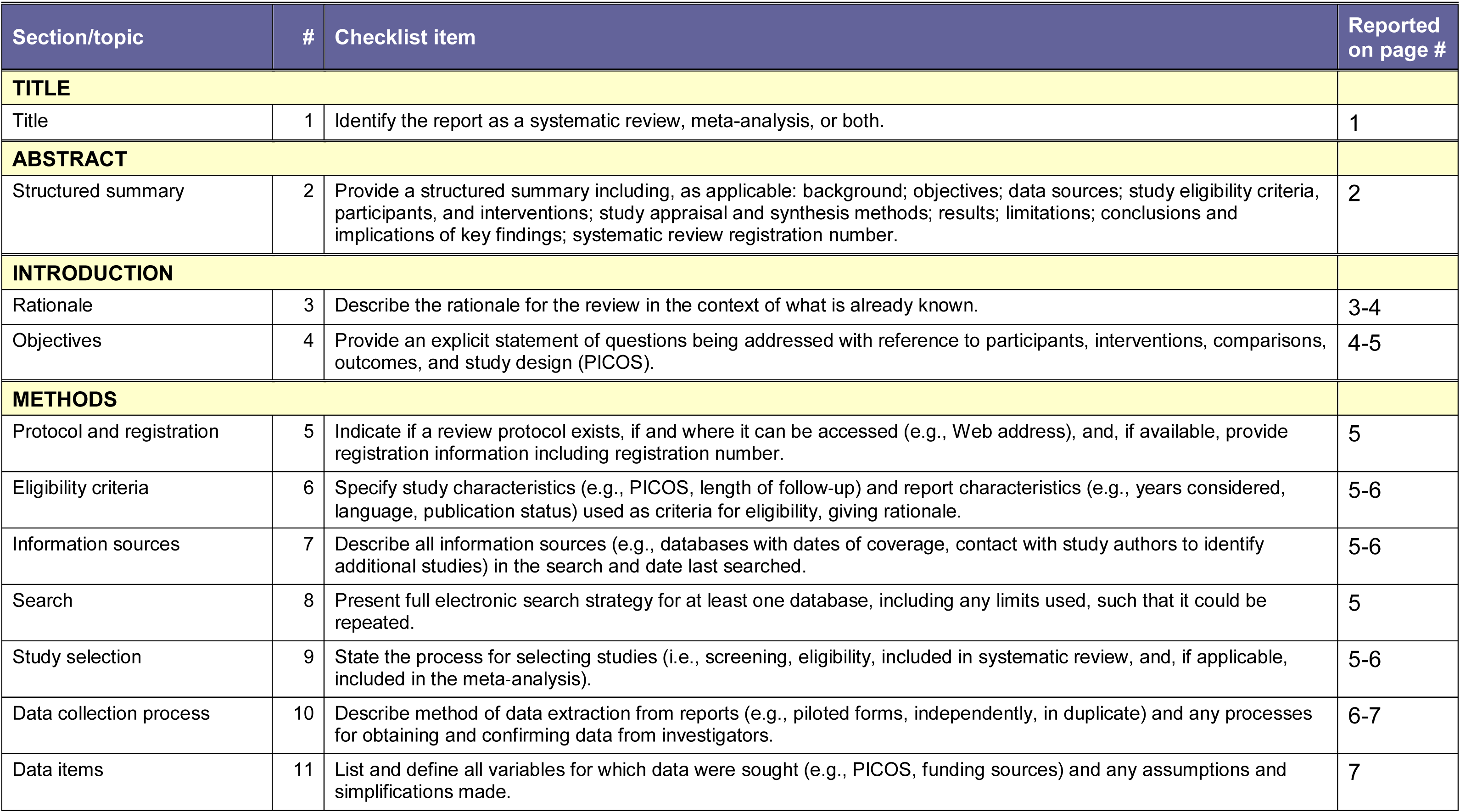
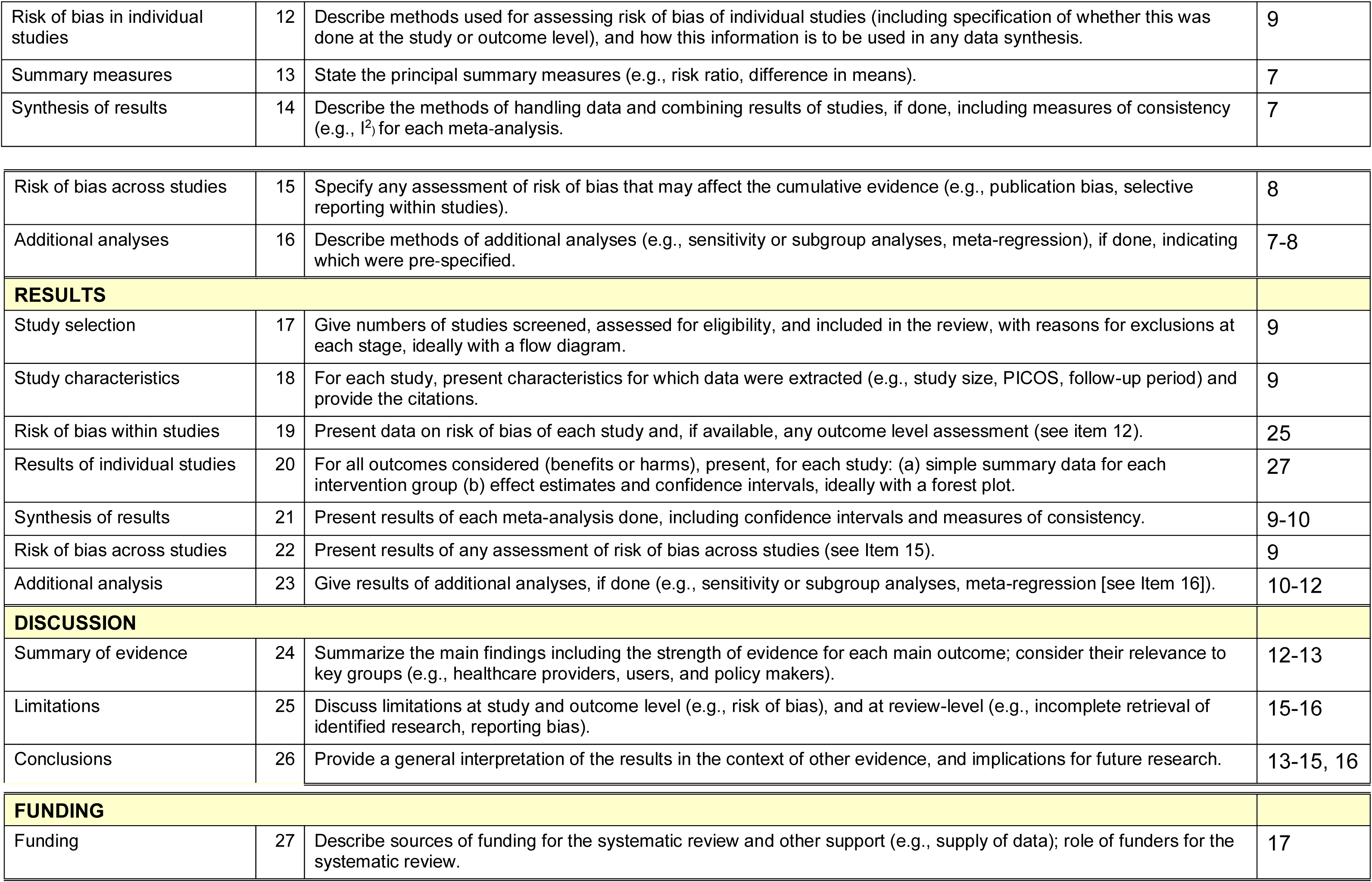

## Supplementary material II: Validation of the web plot digitizer

One method for extracting correlations from articles employed in this meta-analysis was data scraping, in which scatterplots provided in published articles were screenshot and uploaded to a web plot digitizer (22). Three studies (13,55,56) were included because we could scrape data from provided scatterplots. To make sure that this is a precise method for data extraction, we looked at the four studies included in this meta-analysis in which both a correlation and a scatterplot were provided (15-17,53), and compared the correlations obtained from data scraping with the actual correlations stated in the articles. The precision of the web plot digitizer was very high:

| *Study* | *r (web plot digitizer)* | *r (article)* |
| --- | --- | --- |
| *Wang* | 0.922 | 0.926 |
| *Kojima* | 0.597 | 0.596 |
| *Kagerbauer* | -0.121 | -0.149 |
| *Carson* | 0.598 | 0.56 |

There were, however, some slight differences in precision, which may be attributable to differences in scatterplot styles.

## Supplementary material III: Data extraction form for individual studies included in meta-analysis

| Categories | Values |
| --- | --- |
| **A Identification of study** |  |
| 1. Study ID |  |
| 2. Authors |  |
| 3. Title |  |
| 4. Year of publication |  |
| **B Primary values** |  |
| 1. Number of correlations |  |
| 2. Sample size |  |
| 3. Effect size |  |
| 4. Number of samples |  |
| **C Population** |  |
| 1. Species |  |
| 2. Gender |  |
| 3. Mental health status |  |
| 4. Somatic health status |  |
| 5. Age |  |
| **D Moderators** |  |
| 1. Study type |  |
| 2. Sampling location |  |
| 3. Biochemical analysis |  |
| 4. Extraction |  |
| 5. Level of coordination |  |
| **E Other** |  |
| 1. Risk of bias |  |

**A1:** Assign for each study included. **B1:** Specify number of reported correlations. **B2:** For each correlation, specify sample size. **B3:** For each correlation, specify effect size and entity of correlaton. **B4:** For each correlation, specify included number of time-points for coordinated samples. **C3:** Specify mental health status of participants. **C4:** Specify somatic health status of participants. **C5:** Specify age of participants in range and/or mean and standard deviation. **D1:** Specify whether concentrations are sampled at a) baseline level, b) after exogenous OT administration, or c) after other experimental manipulations. **D2:** Specify the locations for central and peripheral samples. **D3:** Specify method for biochemical analysis of OT concentration. **D4:** Specify whether OT was extracted from the sampled substance. **D5:** Specify whether central and peripheral concentrations were sampled a) simultaneously (time interval < 2 mins), b) non-simultaneously, or c) not reported. **E1:** Report risk of bias expressed as a ratio of raw score to obtainable raw score for the study (see risk of bias form).

## Supplementary material IV: Dependent samples variance estimation

For some included studies, multiple effect sizes were reported (e.g. 53). If these effect sizes were independent and representative of different moderators, such as in a between subjects study with an experimental and a control group (e.g. 50), the separate groups were treated as separate studies in the meta-analysis. If, on the other hand, these effect sizes were not independent, such as in studies with repeated measures, preparatory analysis was performed according to planned contingent procedures prior to inclusion of resulting effect sizes to the main meta-study. In the case of repeated measures for the same participants under similar conditions, such as at different time-points in a baseline condition or after exogenous OT administration, separate effect sizes for the different time-points were pooled using a fixed effects model. Variance for the intra-condition pooled effect sizes was estimated with dependent samples variance estimation:

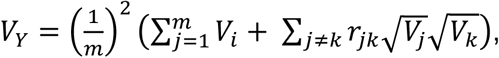

according to the assumption that the intra-condition separate effect sizes were associated (26). In all the cases where exact dependencies were unknown, an r=0.5 association between dependent effect sizes) was assumed, on the premise that the lack of robust variance estimation would default the association on either r=0 or r=1 (26). In the case of repeated measures for the same participants under dissimilar conditions, such as under baseline conditions versus after exogenous OT administration, separate effect sizes expressed different levels of the experimental paradigm moderator variable, and accordingly were not pooled, but rather included in the main meta-analysis directly. However, as these separate effect sizes were not independent, the weight they were afforded in the main meta-analysis was given by dependent samples variance estimation, where a total weight for the pooled effect sizes was distributed to separate effect sizes according to the number of samples in each separate effect size. In other studies, reported single effect sizes were based on repeated measures. For all of these, within-study variance was estimated using dependent samples variance estimation to control for the between measures dependence.

Since exact dependencies between repeated samples were unknown, there is a chance that variances for effect sizes obtained in repeated measures designs were slightly overestimated or underestimated, relative to variances for effect sizes obtained in single sample designs. A differential variance estimation would favor one of the two study types with respect to the relative weight they were afforded in the main analysis. However, since there is no a priori reason to believe that study type should affect effect sizes, this potential bias is not very disquieting. The functions and procedures for variance estimates are available at http://osf/aj55y/

## Supplementary material V: Moderator levels in protocol and in meta-analysis

In the published protocol (21), some of the a priori defined moderator variables were described with other levels than the ones used in the final moderator analysis. Such discrepancies are due to contingent characteristics of included studies that could not be predicted at the stage of meta-analysis planning and protocol registration. For the experimental paradigm moderator, a level was added to accommodate the effect sizes for samples obtained from subjects under experimentally induced stress (47,53,60). In all the included studies, peripheral oxytocin was sampled from blood, such that no levels for peripheral sample location were required. Likewise, sample coordination was high in all included studies, such that no moderator analysis was required for this a priori identified moderator variable.

## Supplementary material VI: Risk of bias tool

**Supplementary material VI: Risk of bias tool.**
| Criteria | Score |
| --- | --- |
| **Category 1: Sample characteristics** |  |
| ***Human population*** |  |
| 1. Presence (or absence) of study-relevant diagnosis established by trained assessor [1p], according to standardized international criteria [1p]. | /2 |
| 2. Reported comorbid diseases [1p] and medication use [1p]. | /2 |
| ***Non-human population*** |  |
| 3. Specified species [1p], source of individual animals stated [1p]. | /2 |
| **Category 2: Internal Validity** |  |
| 4. Potential confounding variables controlled for [2p], or accounted for [1p]. | /2 |
| 5. All [2p] or most [1p] outcome measures have been validated. | /2 |
| **Category 3: Methodology and reporting** |  |
| 6. Method of analysis stated for central [1p] and peripheral [1p] measure. | /2 |
| 7. Outcome data is complete, or incompleteness is accounted for [2p] | /2 |
| 8. Method of extraction described [1p]. | /1 |
| **Category 4: Paradigm-specific criteria** |  |
| 9. IF IN-OT: Proper instructions to participants for administration of OT [1p], measure of nasal cavity health [1p]. | /2 |

For a single study, a raw score of 11-15 is obtainable in principle. For each study, the risk of bias is expressed as a ratio of the actual raw score to its obtainable raw score. Abbreviations: OT = oxytocin, IN-OT = intranasal oxytocin.

## Supplementary material VII: Table of subgroup effects

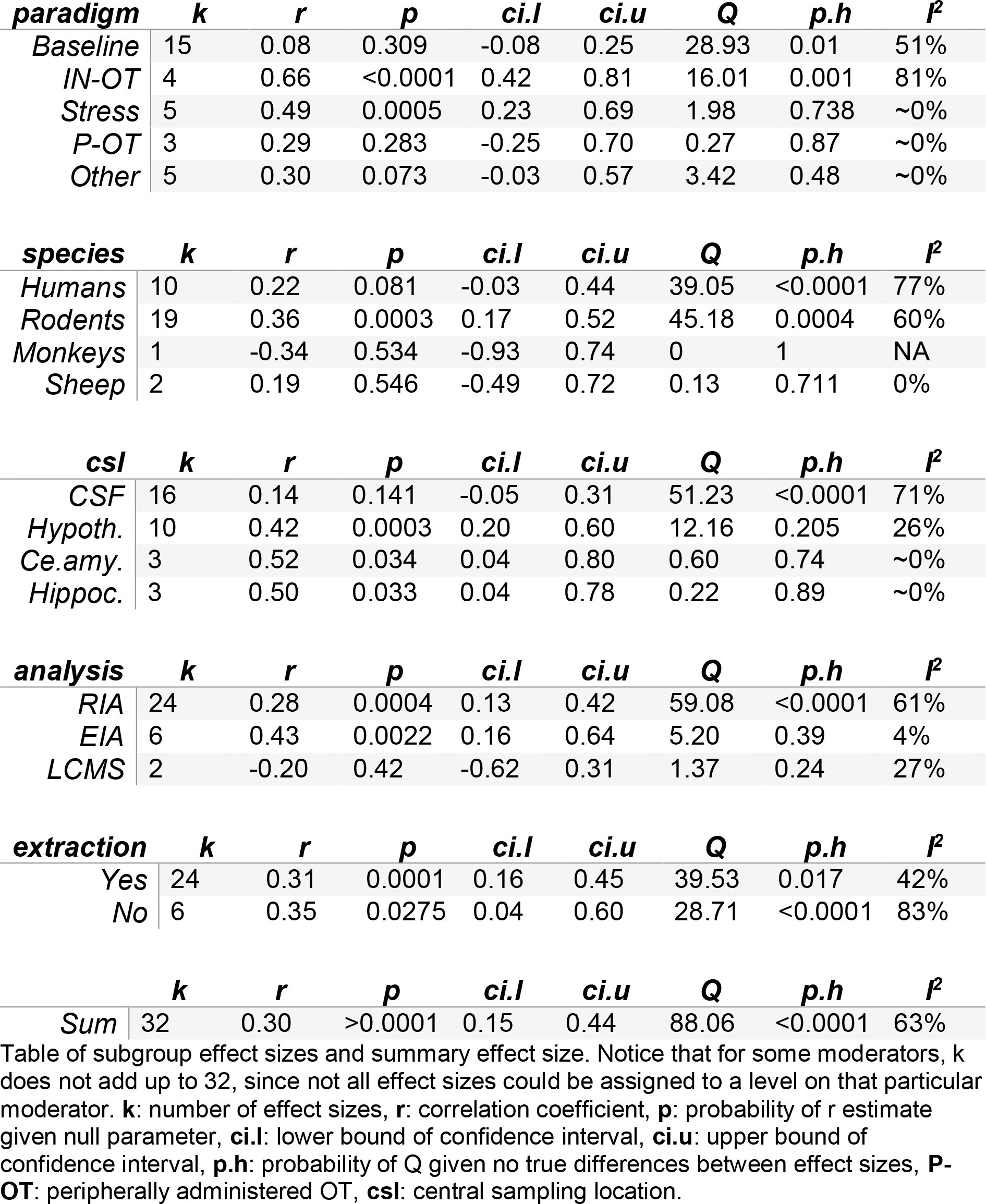

## References

1. Horn J, Swanson L. The autonomic motor system and the hypothalamus. In: Kandel ER, Schwartz JH, Jessel TM, Siegelbaum SA, Hudspeth AJ, editors. Principles of Neural Science. 5 ed. 2013.

2. Neumann ID, Landgraf R. Balance of brain oxytocin and vasopressin: implications for anxiety, depression, and social behaviors. Trends in Neurosciences. 2012 Nov;35(11):649–59.

3. Graustella AJ, MacLeod C. A critical review of the influence of oxytocin nasal spray on social cognition in humans: Evidence and future directions. Hormones and Behavior. 2012 Mar;61(3):410–8.

4. Bartz JA, Zaki J, Bolger N, Ochsner KN. Social effects of oxytocin in humans: context and person matter. Trends in Cognitive Sciences. 2011 Jun.

5. Quintana DS, Guastella AJ, Westlye LT, Andreassen OA. The promise and pitfalls of intranasally administering psychopharmacological agents for the treatment of psychiatric disorders. Molecular Psychiatry. Nature Publishing Group; 2016 Jan 1;21(1):29–38.

6. Guastella AJ, Hickie IB. Oxytocin Treatment, Circuitry, and Autism: A Critical Review of the Literature Placing Oxytocin Into the Autism Context. Biological Psychiatry. 2016 Feb;79(3):234–42.

7. Alvares GA, Quintana DS, Whitehouse AJO. Beyond the hype and hope: Critical considerations for intranasal oxytocin research in autism spectrum disorder. Autism Research. 2016;:1–7.

8. Shilling PD, Feifel D. Potential of Oxytocin in the Treatment of Schizophrenia. CNS Drugs. Springer International Publishing; 2016;30(3):193–208.

9. McCullough ME, Churchland PS, Mendez AJ. Problems with measuring peripheral oxytocin: Can the data on oxytocin and human behavior be trusted? Neuroscience & Biobehavioral Reviews. 2013 Sep;37(8):1485–92.

10. Wotjak CT, Ganster J, Kohl G, Holsboer F, Landgraf R, Engelmann M. Dissociated central and peripheral release of vasopressin, but not oxytocin, in response to repeated swim stress: New insights into the secretory capacities of peptidergic neurons. Neuroscience. 1998 Aug;85(4):1209–22.

11. Ross HE, Cole CD, Smith Y, Neumann ID, Landgraf R, Murphy AZ, et al. Characterization of the oxytocin system regulating affiliative behavior in female prairie voles. Neuroscience. 2009 Sep;162(4):892–903.

12. Landgraf R, Neumann I, Schwarzberg H. Central and peripheral release of vasopressin and oxytocin in the conscious rat after osmotic stimulation. Brain Research. 1988 Aug;457(2):219–25.

13. Amico JA, Challinor SM, Cameron JL. Pattern of Oxytocin Concentrations in the Plasma and Cerebrospinal Fluid of Lactating Rhesus Monkeys (Macaca mulatto,): Evidence for Functionally Independent Oxytocinergic Pathways in Primates*. The Journal of Clinical Endocrinology & Metabolism. The Endocrine Society; 1990 Jul 1;71(6): 1531–5.

14. Robinson ICAF, Jones PM. Oxytocin and Neurophysin in Plasma and CSF during Suckling in the Guinea-Pig. Neuroendocrinology. Karger Publishers; 1982 Jul 1;34(1):59–63.

15. Carson DS, Berquist SW, Trujillo TH, Garner JP, Hannah SL, Hyde SA, et al. Cerebrospinal fluid and plasma oxytocin concentrations are positively correlated and negatively predict anxiety in children. Molecular Psychiatry. Nature Publishing Group; 2014 Nov 4;20(9):1085–90.

16. Kagerbauer SM, Martin J, Schuster T, Blobner M, Kochs EF, Landgraf R. Plasma Oxytocin and Vasopressin do not Predict Neuropeptide Concentrations in Human Cerebrospinal Fluid. Journal of Neuroendocrinology. 2013 Jul 1;25(7):668–73.

17. Wang Y-L, Yuan Y, Yang J, Wang C-H, Pan Y-J, Lu L, et al. The interaction between the oxytocin and pain modulation in headache patients. Neuropeptides. 2013 Apr;47(2):93–7.

18. Striepens N, Kendrick KM, Hanking V, Landgraf R, Wüllner U, Maier W, et al. Elevated cerebrospinal fluid and blood concentrations of oxytocin following its intranasal administration in humans. Scientific Reports. Nature Publishing Group; 2013 Dec 6;3:3440.

19. Moher D, Liberati A, Tetzlaff J, Altman DG. Preferred Reporting Items for Systematic Reviews and Meta-Analyses: The PRISMA Statement. Ann Intern Med. American College of Physicians; 2009 Aug 18;151(4):264–9.

20. Quintana DS. From pre-registration to publication: a non-technical primer for conducting a meta-analysis to synthesize correlational data. Front Psychol. Frontiers Media SA; 2015;6(80):839.

21. Valstad M, Alvares GA, Andreassen OA, Westlye LT, Quintana DS. The relationship between central and peripheral oxytocin concentrations: a systematic review and meta-analysis protocol. Systematic Reviews 2016 5:1. BioMed Central; 2016 Mar 31;5(1):1.

22. Rohatgi A. WebPlotDigitizer [Internet]. 3rd ed. Austin, Texas. Available from: http://arohatgi.info/WebPlotDigitizer/

23. Del Re AC, Hoyt WT. MAc: Meta-analysis with correlations. 1st ed. Available from: https://CRAN.R-project/package=MAc

24. Viechtbauer W. Conducting meta-analyses in R with the metafor package. J Stat Softw. 2010.

25. Hothorn T, Bretz F, Westfall P. Simultaneous Inference in General Parametric Models. Biometrical Journal. WILEY-VCH Verlag; 2008 Jun 1; 50(3):346–63.

26. Borenstein MH, Higgins LV, Rothstein J. Introduction to meta-analysis. Chichester, England: Wiley; 2009.

27. Gilpin AR. Table for Conversion of Kendall“S Tau to Spearman”S Rho Within the Context of Measures of Magnitude of Effect for Meta-Analysis. Educational and Psychological Measurement. SAGE Publications; 1993 Mar 1; 53(1):87–92.

28. DerSimonian R, Kacker R. Random-effects model for meta-analysis of clinical trials: An update. Contemporary Clinical Trials. Elsevier; 2007 Feb 1; 28(2):105–14.

29. Higgins JPT, Thompson SG, Deeks JJ, Altman DG. Measuring inconsistency in meta-analyses. BMJ. British Medical Journal Publishing Group; 2003 Sep 4;327(7414):557–60.

30. Egger M, Smith GD, Schneider M, Minder C. Bias in meta-analysis detected by a simple, graphical test. BMJ. British Medical Journal Publishing Group; 1997 Sep 13;315(7109):629–34.

31. Schulz KF, Chalmers I, Hayes RJ, Altman DG. Empirical Evidence of Bias: Dimensions of Methodological Quality Associated With Estimates of Treatment Effects in Controlled Trials. JAMA. American Medical Association; 1995 Feb 1;273(5):408–12.

32. Peters JL, Sutton AJ, Jones DR, Abrams KR, Rushton L. Contour-enhanced meta-analysis funnel plots help distinguish publication bias from other causes of asymmetry. Journal of Clinical Epidemiology. 2008 Oct;61(10):991–6.

33. Alvares GA, Quintana DS, Hickie IB, Guastella AJ. Autonomic nervous system dysfunction in psychiatric disorders and the impact of psychotropic medications: a systematic review and meta-analysis. J. Psychiatry Neurosci; 2015.

34. Hoge EA, Pollack MH, Kaufman RE, Zak PJ, Simon NM. Oxytocin Levels in Social Anxiety Disorder. CNS Neuroscience & Therapeutics. Blackwell Publishing Inc; 2008 Sep 1;14(3):165–70.

35. Rubin LH, Carter CS, Drogos L, Pournajafi-Nazarloo H, Sweeney JA, Maki PM. Peripheral oxytocin is associated with reduced symptom severity in schizophrenia. Schizophrenia Research. 2010 Dec;124(1-3):13–21.

36. Lane A, Luminet O, Nave G, Mikolajczak M. Is there a publication bias in behavioral intranasal oxytocin research on humans? Opening the file drawer of one lab. Journal of Neuroendocrinology. 2016.

37. Walum H, Waldman ID, Young LJ. Statistical and Methodological Considerations for the Interpretation of Intranasal Oxytocin Studies. Biological Psychiatry. 2016 Feb;79(3):251–7.

38. Guastella AJ, Mitchell PB, Dadds MR. Oxytocin Increases Gaze to the Eye Region of Human Faces. Biological Psychiatry. 2008 Jan;63(1):3–5.

39. Andari E, Duhamel J-R, Zalla T, Herbrecht E, Leboyer M, Sirigu A. Promoting social behavior with oxytocin in high-functioning autism spectrum disorders. PNAS. National Acad Sciences; 2010 Mar 2;107(9):4389–94.

40. Domes G, Heinrichs M, Gläscher J, Büchel C, Braus DF, Herpertz SC. Oxytocin Attenuates Amygdala Responses to Emotional Faces Regardless of Valence. Biological Psychiatry. 2007 Nov;62(10):1187–90.

41. Kosfeld M, Heinrichs M, Zak PJ, Fischbacher U, Fehr E. Oxytocin increases trust in humans. Nature. Nature Publishing Group; 2005 Jun 2;435(7042):673–6.

42. Born J. Sniffing neuropeptides: a transnasal approach to the human brain. Nat Neurosci. 2002;5:514–516PB–M3–N1–UR–.

43. Quintana DS, Alvares GA, Hickie IB, Guastella AJ. Do delivery routes of intranasally administered oxytocin account for observed effects on social cognition and behavior? A two-level model. Neuroscience & Biobehavioral Reviews. 2015 Feb;49:182–92.

44. Quintana DS, Westlye LT, Rustan ØG, Tesli N, Poppy CL, Smevik H, et al. Low-dose oxytocin delivered intranasally with Breath Powered device affects social-cognitive behavior: a randomized four-way crossover trial with nasal cavity dimension assessment. Transl Psychiatry. 2014 Dec 1; 5(7):e602–2.

45. Quintana DS, Westlye LT, Alnæs D, Rustan ØG, Kaufmann T, Smerud KT, et al. Low dose intranasal oxytocin delivered with Breath Powered device dampens amygdala response to emotional stimuli: A peripheral effect-controlled within-subjects randomized dose-response fMRI trial. Psychoneuroendocrinology. 2016 Jul;69:180–8.

46. Kojima S, Stewart RA, Demas GE, Alberts JR. Maternal Contact Differentially Modulates Central and Peripheral Oxytocin in Rat Pups During a Brief Regime of Mother–Pup Interaction that Induces a Filial Huddling Preference. Journal of Neuroendocrinology. Blackwell Publishing Ltd; 2012 May 1;24(5):831–40.

47. Williams SK, Barber JS, Jamieson Drake AW, Enns JA, Townsend LB, Walker CH, et al. Chronic Cocaine Exposure During Pregnancy Increases Postpartum Neuroendocrine Stress Responses. Journal of Neuroendocrinology. Blackwell Publishing Ltd; 2012 Apr 1; 24(4):701–11

48. Ferguson AV, Latchford KJ, Samson WK. The paraventricular nucleus of the hypothalamus – a potential target for integrative treatment of autonomic dysfunction. Expert Opinion on Therapeutic Targets. Taylor & Francis; 2008 May 15;12(6):717–27.

49. Yang J, Yang Y, Chen J-M, Liu W-Y, Wang C-H, Lin B-C. Central oxytocin enhances antinociception in the rat. Peptides. 2007 May;28(5):1113–9.

50. Martin J, Kagerbauer SM, Schuster T, Blobner M, Kochs EF, Landgraf R. Vasopressin and oxytocin in CSF and plasma of patients with aneurysmal subarachnoid haemorrhage. Neuropeptides. 2014 Apr;48(2):91–6.

51. Kagerbauer SM. Plasma oxytocin and vasopressin do not predict neuropeptide concentrations in the human cerebrospinal fluid. J Neuroendocrinol. 2013;25:668–673PB–M3–N1–UR–.

52. Neumann ID, Maloumby R, Beiderbeck DI, Lukas M, Landgraf R. Increased brain and plasma oxytocin after nasal and peripheral administration in rats and mice. Psychoneuroendocrinology. 2013 Oct;38(10): 1985–93.

53. Kojima S, Stewart RA, Demas GE, Alberts JR. Maternal Contact Differentially Modulates Central and Peripheral Oxytocin in Rat Pups During a Brief Regime of Mother–Pup Interaction that Induces a Filial Huddling Preference. Journal of Neuroendocrinology. Blackwell Publishing Ltd; 2012 May 1;24(5):831–40.

54. Sansone GR, Gerdes CA, Steinman JL, Winslow JT, Ottenweller JE, Komisaruk BR, et al. Vaginocervical Stimulation Releases Oxytocin within the Spinal Cord in Rats. Neuroendocrinology. Karger Publishers; 2002 Apr 29;75(5):306–15.

55. Takeda S, Kuwabara Y, Mizuno M. Effects of pregnancy and labor on oxytocin levels in human plasma and cerebrospinal fluid. Endocrinologia Japonica. 1985;32(6):875–80.

56. Takagi T, Tanizawa O, Otsuki Y, Sugita N, Haruta M, Yamaji K. Oxytocin in the cerebrospinal fluid and plasma of pregnant and nonpregnant subjects. Horm Metab Res. 1985 Jun 1;17(6):308–10.

57. Jokinen J, Chatzittofis A, Hellström C, Nordström P, Uvnäs-Moberg K, Åsberg M. Low CSF oxytocin reflects high intent in suicide attempters. Psychoneuroendocrinology. 2012 Apr;37(4):482–90.

58. Jin D, Liu H-X, Hirai H, Torashima T, Nagai T, Lopatina O, et al. CD38 is critical for social behaviour by regulating oxytocin secretion. Nature. Nature Publishing Group; 2007 Mar 1;446(7131):41–5.

59. Keverne EB, Kendrick KM. Morphine and corticotrophin-releasing factor potentiate maternal acceptance in multiparous ewes after vaginocervical stimulation. Brain Research. 1991 Feb;540(1-2):55–62.

60. Engelmann M, Bull PM, Brown CH, Landgraf R, Horn TFW, Singewald N, et al. GABA selectively controls the secretory activity of oxytocin neurons in the rat supraoptic nucleus. European Journal of Neuroscience. Blackwell Science Ltd; 2004 Feb 1;19(3):601–8.

61. Kleindienst A, Hildebrandt G, Kroemer SA, Franke G, Gaab MR, Landgraf R. Hypothalamic neuropeptide release after experimental subarachnoid hemorrhage: in vivo microdialysis study. Acta Neurologica Scandinavica. Blackwell Publishing; 2004 May 1;109(5):361–8.

